# Face in the facade: How face-likeness modulates memory and neural representations

**DOI:** 10.64898/2026.05.06.723204

**Authors:** Claire Pauley, Izabela Maria Sztuka, Nour Tawil, Simone Kühn

## Abstract

Evidence suggests that information represented more reliably in neural activity patterns across repeated exposures is more likely to be remembered. However, this relationship varies across category-selective regions of the ventral visual cortex. Specifically, for house stimuli neural reliability has been robustly linked to memory outcomes in the parahippocampal place area (PPA), but less consistently for faces in the fusiform face area (FFA). The reason for this mismatch is unknown. To address this discrepancy, we implemented a novel within-category manipulation by presenting highly face-like and non-face-like house stimuli during fMRI, followed by a memory test. Non-face-like houses were more likely to be remembered than face-like houses. Although face-likeness did not elicit face-selective responses in the FFA, representational reliability in ventral visual cortices, particularly in the FFA, showed an association with individual differences in memory performance. Finally, symmetry emerged as a potential perceptual factor underlying differences in mnemonic outcomes.

## 1 Introduction

Successful remembering relies on the formation of stable memory representations that reliably reflect event-related information, and distinguish a particular event from other, similar events. Accordingly, greater consistency in the representation of individual stimuli in neural activity patterns across repeated presentations has been shown to predict successful memory of these stimuli (see Figure 1 for schematic; Kuhl & Chun, 2014; Xue et al., 2010).

**Figure 1.**
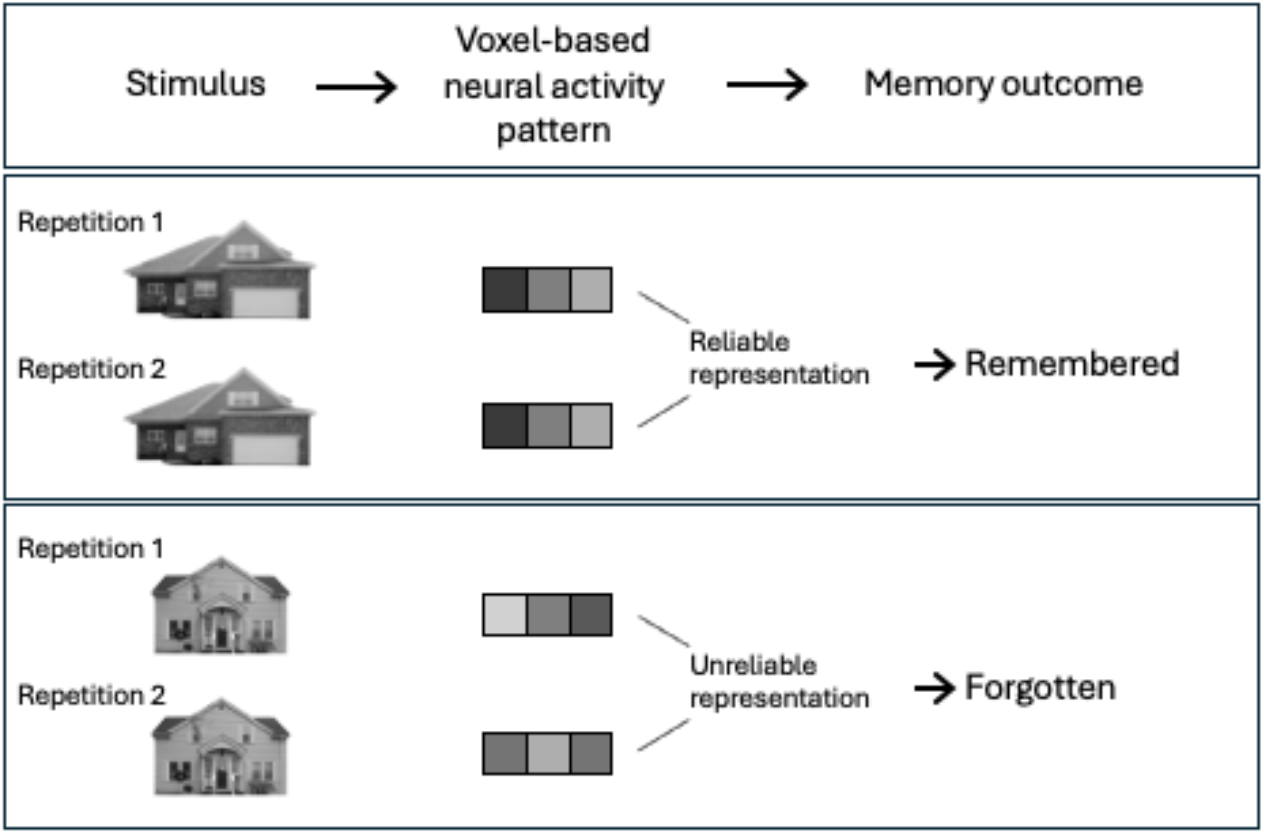
Simplified schematic depicting greater probability of remembering stimuli with reliable neural representations across repetitions (middle panel). Stimuli with unreliable neural representations are less likely to be remembered (lower panel). Neural representations are portrayed as different shades of grey representing voxel-based neural activity patterns (3 voxel example).

Memory studies evaluating the degree to which stimuli are distinguishable from one another based on their neural representations often exploit the specialized categorical processing of the ventral temporal cortices. Specifically, parts of the fusiform gyrus (fusiform face area; FFA) are selectively involved in processing face stimuli (Kanwisher et al., 1997), while the parahippocampal gyrus (parahippocampal place area; PPA) selectively processes house and scene stimuli (Epstein & Kanwisher, 1998). Evidence linking neural distinctiveness to memory outcomes ostensibly varies between the FFA and PPA. For example, Hasinski and Sederberg (2016) found that face-specific pattern similarity in the FFA was greater for faces that were subsequently remembered rather than forgotten (see also, Xue et al., 2010). However, such fine-grained associations between memory outcomes and neural representations in the FFA are not consistently identified (e.g., Pauley et al., 2023). Turning to the PPA, findings more consistently demonstrate the role of highly specific neural representations of scene and house stimuli to support trial-by-trial memory outcomes (Pauley et al., 2023), as well as explaining interindividual differences in memory performance (Koen et al., 2019; Pauley et al., 2023; Srokova et al., 2020).

Thus, prior evidence suggests the link between distinctive neural representations and memory outcomes may be more consistent for house/scene processing in the PPA relative to face processing in the FFA. However, the origins of this incongruity are unclear. This difference may stem from regional (e.g., morphological) variations between the PPA and FFA or from stimulus-specific differences between house and face stimuli. For example, stimulus properties, including stimulus complexity (Garrett et al., 2020), inherently differ between houses and faces. Faces have clear structures (e.g., two eyes positioned above a nose and a mouth), which reduces the perceptual dimensionality varying between individual exemplars. In contrast, houses have far fewer structural constraints, resulting in substantial variation between individual house exemplars. Disentangling the factors contributing to processing differences between the PPA and FFA proves a challenge due to the fact that region and stimulus are inextricably intertwined (the PPA preferentially processes houses, while the FFA preferentially processes faces).

The human tendency to anthropomorphize non-human objects could provide an opportunity to disentangle the role of stimulus-specific processing differences between the FFA and PPA. People recognize human-like attributes in diverse objects (Guthrie, 1995), including cars, robots, and houses (see also, Pohlmann et al., 2024). Such anthropomorphic tendencies are further traceable in neural activity. Face-like activation has been recorded in the FFA in response to non-face stimuli when these stimuli have face-like perceptual properties. Specifically, stronger anthropomorphism of cars has been associated with greater activation in the FFA (Kühn et al., 2014), demonstrating that also non-face stimuli can elicit face-like neural signatures.

In this study, we seek to understand how neural representations of houses in the PPA and FFA track memory outcomes. We presented images of individual house facades, allowing us to manipulate and measure stimulus-specific properties. Crucially, half of the house stimuli were independently rated as highly face-like, while the other half were rated very low on face-likeness (Filliter et al., 2016). We predicted that face-like houses would elicit face-like activation in the FFA and less face-like houses would elicit house-like activation in the PPA. We predicted that this stimulus-selective response pattern would allow us to isolate regional differences in the relationship between neural reliability and memory outcomes with shared stimulus properties across conditions.

## 2 Materials and Methods

The present study questions and analyses were pre-registered before the start of any data analysis (https://aspredicted.org/wqdb-tgjv.pdf).

### 2.1 Participants

Twenty-five participants took part in the current experiment (11 females, 14 males). Participants were 18-48 years of age (mean = 32.6 years, SD = 8.9 years) and had no history of psychiatric or neurological disorders. Twenty additional participants were collected but were excluded from data analyses: 11 were administered a different version of the memory test (which was altered after memory performance rarely exceeded chance), 1 participated twice, 1 repeated one of the scanner runs, for 2 data was lost due to technical errors, and 5 performed below chance on the memory test. The sample size was initially determined using a power analysis and adding in several additional participants in case some were lost due to poor data quality. Although more were excluded from the present analysis than expected due to the change in memory paradigm, the present sample size still aligns with recent fMRI studies assessing multivariate memory representations (e.g., Howard et al., 2024). The study was approved by the local psychological ethics committee of the University Medical Center Hamburg-Eppendorf (LPEK-0697) and written informed consent was obtained from each participant prior to study participation.

### 2.2 Paradigm

Participants completed a localizer task and a main houses task inside the MRI scanner. For the localizer task, blocks of grayscale face and house stimuli (stimuli obtained from Haxby et al., 2001) were presented for 500ms per stimulus separated by a white fixation cross (1s). Eight blocks (4 face blocks, 4 house blocks) of 12 stimuli each were presented with a 10s fixation cross between each block. Participants were instructed to keep their eyes open and focus on the stimuli. For the main houses task, participants completed two identical runs in which 40 images of houses were presented 3 times each (1 s duration) interleaved with jittered fixation crosses (2.5 – 5.5 s). The house images were obtained from the DalHouses stimulus set (Filliter et al., 2016), in which 100 grayscale houses were subjectively rated on several measures including face-likeness. Based on these independent ratings, we selected the 20 highest- and lowest-rated houses (termed here highFL and lowFL, respectively; see Figure 2). Due to the limited size of the original images, we upscaled them using PixelCut’s AI-based Image Upscaler (https://www.pixelcut.ai/image-upscaler), doubling their resolution. Minor corrections were applied in Adobe Photoshop (Adobe Inc., 2024) to restore architectural details not fully preserved by the automated process. Participants were instructed to press a button when the fixation cross turned red (3 times per run) to keep them attentive throughout the session. Following MRI scanning, participants completed two surprise memory tests: one old/new recognition test and one forced-choice memory test. For the old/new recognition test, the next 20 highest- and lowest-rated face-like houses were selected to serve as the “new” never-before-seen images. On each trial (80 in total), a house image was displayed for 3 s and participants were asked to indicate as quickly as possible whether they had seen the image before (“old”) or whether they had never seen the image before (“new”). A fixation cross was displayed for 2 s between each trial. For the forced-choice recognition test, the 10 highest- and lowest-rated face-like houses were edited using Adobe Photoshop with minor feature changes (e.g., in the details of windows, doors, or railings) to create a set of highly similar house images. Each house was presented side-by-side with its altered pair and participants were instructed to indicate which house they have already seen before using the left and right arrow keys (20 trials in total). The stimuli were displayed for 4 s while participants made the memory judgment, and each trial was separated by a 500 ms fixation cross. Following the memory tests, participants were asked to rate each of the “old” houses on 6 dimensions, including face-likeness (very much-not at all), preference (very much-not at all), naturalness (very natural-very artificial), safety (approach-avoid), arousal (very stressful-not at all stressful), and relaxing (very much-not at all). Each dimension was rated on an analogue sliding scale with possible responses between 0 and 100.

**Figure 2.**
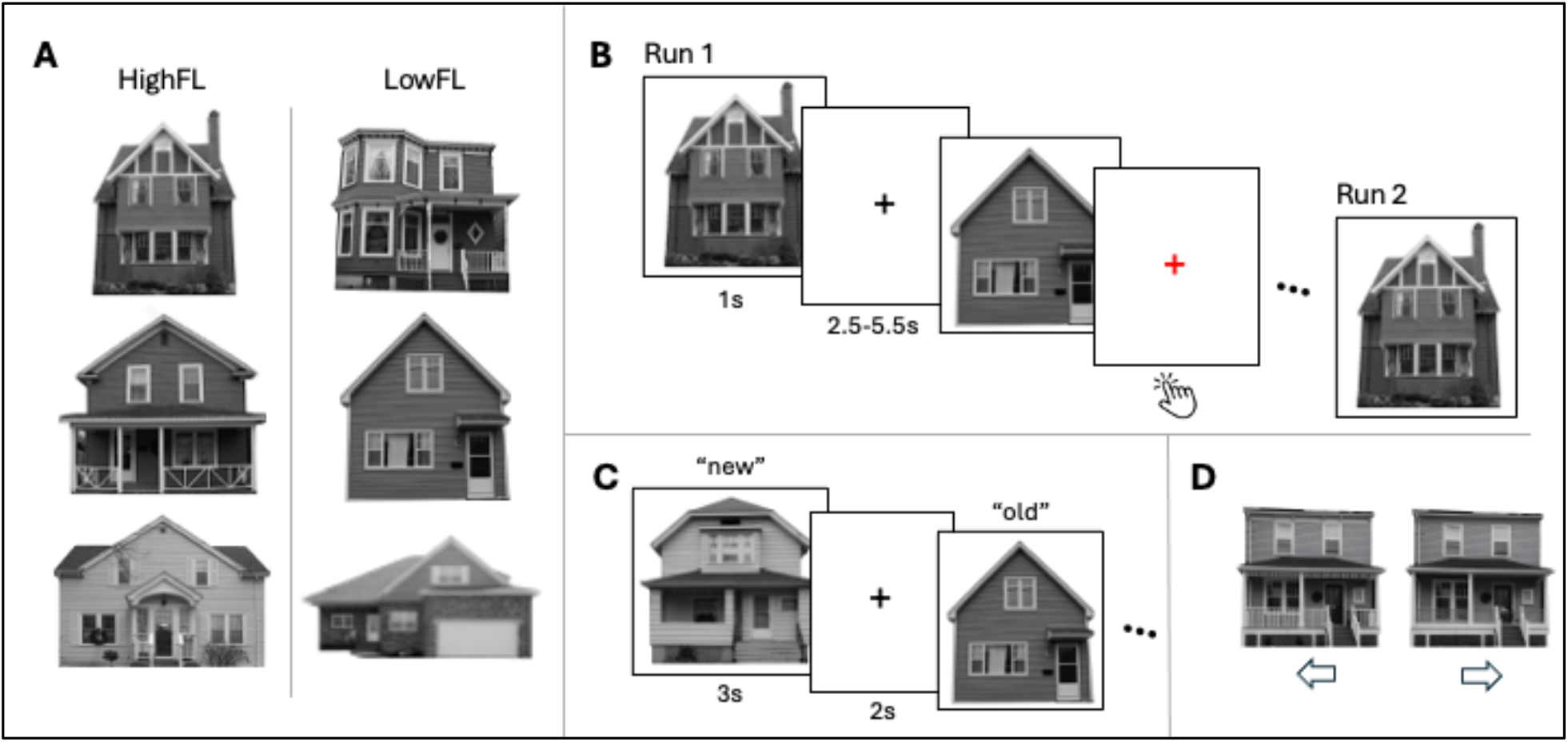
Experimental design. (A) Example stimuli from the high face-like (highFL) and low face-like (lowFL) stimulus conditions. (B) During the main task, houses were presented one-by-one with a jittered interstimulus interval. Each run consisted of 3 blocks in which each house was presented once and both runs had identical presentation order. Participants completed a target detection task throughout, in which they pressed a button when the fixation cross turned red. (C) After scanning, participants completed a surprise memory test in which they indicated whether they had previously seen the displayed house (“old”) or whether they had not seen the house before (“new”). (D) Participants additionally completed a forced-choice memory test in which “old” house facades were slightly altered and participants were asked to indicate which house (left or right) they had previously seen.

### 2.3 Behavioral analyses

Behavioral analyses were performed using custom MATLAB scripts. We first evaluated how well the classification of highFL and lowFL (based on the independent ratings in Filliter et al., 2016) aligned with the subjective ratings of face-likeness in the current sample. For each participant, a median split divided the 40 images from the main houses task based on each individuals’ subjective ratings of face-likeness. Images with median ratings or below were marked as lowFL and images rated higher than the median were marked as highFL. We report the percentage of subjective classifications aligning with the independent ratings.

We were then interested in whether face-likeness modulated memory performance. Recognition memory performance (*Pr*) was calculated as the difference between the hit rate (proportion of correctly identified “old” houses) and the false alarm rate (proportion of “new” houses incorrectly identified as old; Snodgrass & Corwin, 1988). Participants with below-chance memory performance (*Pr* < 0) were excluded from further analyses (see also Section 2.1). Forced-choice memory performance was measured as the proportion of correctly-identified “old” houses. Dependent-samples *t*-tests were conducted to determine if memory performance differed between highFL and lowFL houses. The interindividual relationship between *Pr* and forced-choice memory performance was assessed using Pearson correlation.

### 2.4 fMRI data acquisition and preprocessing

MRI scanning was performed using a Siemens Magnetom TimTrio 3T MRI scanner with a 32-channel head-coil at the Max Planck Institute for Human Development in Berlin, Germany. First, a T1-weighted (T1w) magnetization prepared rapid acquisition gradient echo (MPRAGE) pulse sequence image was acquired (voxel size = 1 x 1 x 1 mm^3^; TR = 2.5 s; TE = 4.77 ms; flip angle = 7◦; TI = 1.1 s). Functional images for the localizer and main houses tasks were collected using a whole-brain multi-band gradient echo planar imaging (EPI) sequence (multiband acceleration factor = 2; voxel size: 2 × 2 × 2 mm^3^; TR = 2 s; TE = 29.6 ms; flip angle = 80°; FoV = 192 mm). Fieldmaps were collected for signal distortion correction (TR = 2.25 s; TE = 38.8 ms; flip angle = 90°). A functional resting state scan was additionally acquired, but not used in the present analyses.

MRI data were preprocessed using *fMRIPrep* (version 23.2.3; Esteban et al., 2019) with the default settings. The T1w image was corrected for intensity nonuniformity, skull-stripped, and normalized to the ICBM 152 Nonlinear Asymmetrical template version 2009c through nonlinear registration. Functional images were motion-corrected, slice-time corrected, corrected for field distortion, and co-registered to the normalized T1w reference image. Functional images for the localizer task were smoothed using an 8mm FWHM Gaussian kernel and functional images for the main houses task were smoothed using a 2mm FWHM Gaussian kernel. All analyses were conducted in MNI space.

### 2.5 Participant-specific FFA and PPA ROIs

The functional localizer run was used to isolate brain regions selectively responsive to face and house stimuli. We implemented a GLM with a block design, including a face regressor, a house regressor, and six motion parameters as nuisance regressors. Based on the preferential processing of face and house stimuli in the fusiform gyrus and parahippocampal gyrus, respectively (Epstein & Kanwisher, 1998; Kanwisher et al., 1997), we considered these regions as defined by the AAL3 atlas (Tzourio-Mazoyer et al., 2002). To create participant-specific localizer maps, the voxels in the fusiform gyrus were sorted from the highest to lowest *t*-values for the contrast faces > houses independently for each participant. Comparably, in the parahippocampal gyrus, the voxels were sorted from the highest to lowest *t*-values for the houses > faces contrast for each participant. The top 250 and 500 voxels in each region were isolated, resulting in two unique FFA and PPA ROIs for each individual (see Figure 3). We additionally evaluated template ROIs in which all voxels of the FFA and PPA were included.

**Figure 3.**
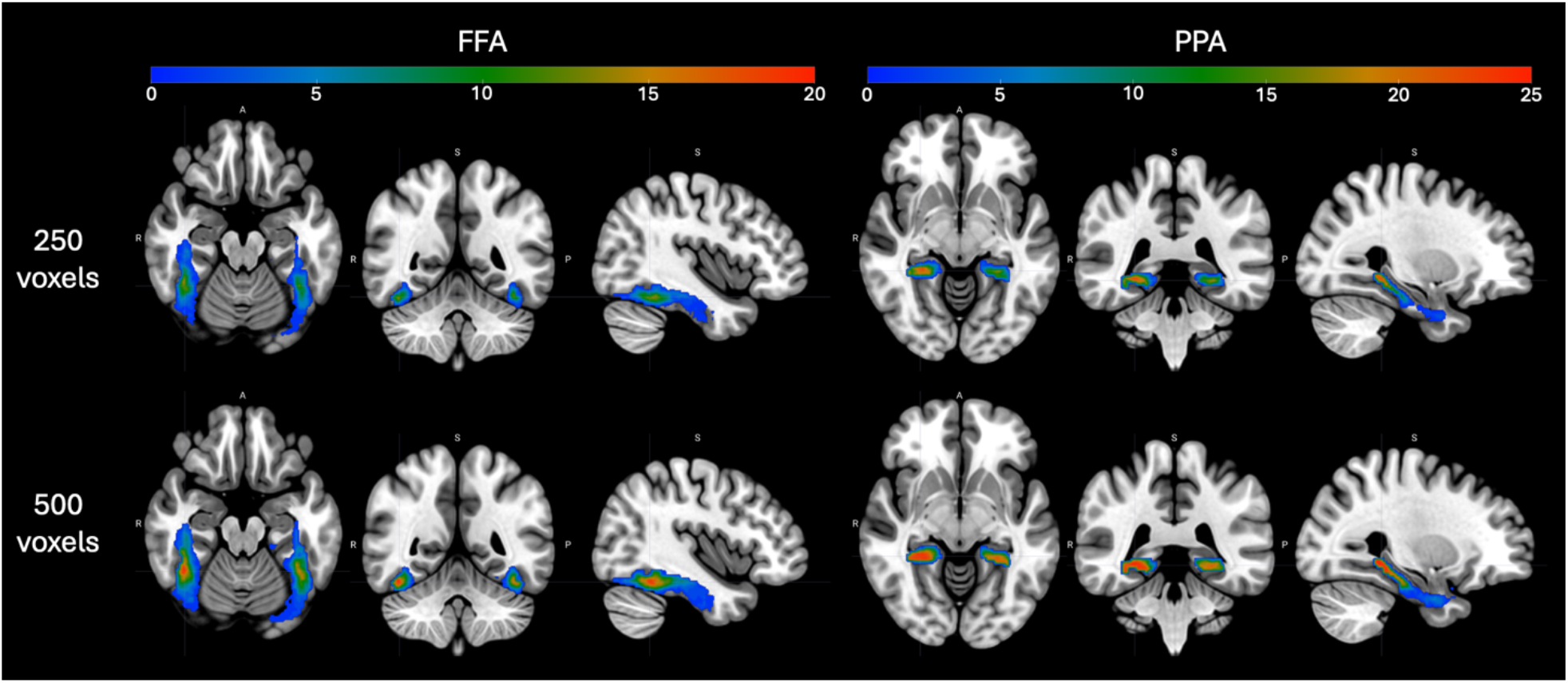
Probabilistic maps showing the distribution of voxels isolated in the localizer analysis. Individual voxels are colored based on the number of participants in which a voxel was localized (maximum 25; note that the scales differ between the FFA, left, and PPA, right).

### 2.6 Assessing neural differences between highFL and lowFL houses

We were interested in evaluating the reliability of neural representations across repeated presentations of the same stimuli. To this end, we used pattern similarity analysis to assess stimulus-specific neural responses to highFL and lowFL houses. Single-trial GLMs were conducted for each individual trial using a Least Squares Separate (LSS) design (Mumford et al., 2012) with one trial-specific regressor, one regressor modeling all other trials in the run, and six motion parameters as nuisance regressors. Trials were modeled as event-related stick functions corresponding with the onset of each stimulus convolved with a canonical hemodynamic response function. Pattern similarity analyses were conducted on the resulting β weights for each trial using Pearson correlations. Only correlations between trials from different runs were considered to control for autocorrelations in the hemodynamic responses (Dimsdale-Zucker & Ranganath, 2018). Additionally, trials were matched based on order of presentation within each run to control for repetition-related artifacts in the neural activity patterns (Grill-Spector et al., 2006). Stimulus-specific reliability was modeled as the interaction between within-stimulus similarity and between-stimulus similarity (see Pauley et al., 2023, 2024, for similar methodology). Within-stimulus similarity was defined as the mean of the correlations of across-voxel activation patterns of the same stimulus to itself across repetitions. Between-stimulus similarity was defined as the mean of the correlations of a particular stimulus to all other stimuli of the same face-likeness categorization. To investigate how face-likeness modulated neural representations in the FFA and PPA, stimulus-specific reliability was tested using 2 (within-stimulus/between-stimulus) x 2 (highFL/lowFL) x 2 (FFA/PPA) ANOVAs. Follow-up 2×2 ANOVAs were conducted within each ROI independently. ANOVAs were assessed for both levels of the participant-specific ROIs (top 250 and top 500 selective voxels in the fusiform and parahippocampal gyri) as well as for all voxels in these regions. Dependent-samples *t*-tests were conducted to test whether reliability differed significantly from zero.

### 2.7 Whole-brain searchlight similarity analyses

Searchlight similarity analyses were implemented using customized scripts based on the MATLAB toolbox for representational similarity analysis (rsatoolbox 2.0; Nili et al., 2014) using 8-mm-radius spherical searchlights. Stimulus-specific reliability (as defined in Section 2.6) was calculated in a searchlight centered on each voxel, resulting whole-brain maps of representational reliability for high- and lowFL houses. Thresholding was implemented using nonparametric, cluster-based, random permutation analyses adapted from the FieldTrip toolbox (Oostenveld et al., 2011) to identify brain regions demonstrating differences in neural reliability between high- and lowFL houses. First, dependent-samples *t*-tests were conducted in each voxel. Adjacent voxels below a threshold of *p* < 0.005 were grouped together into clusters. The sum of all *t* statistics in each cluster was defined as the cluster test statistic. The Monte Carlo method was used to determine whether a cluster was significant by comparing the cluster test statistic to a reference distribution of *t* statistics across 1000 permutations. Each *t* statistic in the reference distribution was created by randomly reallocating the two conditions and calculating the cluster test statistic based on this random reallocation. Clusters were considered significant under a threshold of *p* < 0.05 and if they contained at least 10 voxels.

### 2.8 Exploring the relationship between neural reliability and memory performance

In order to determine the possible influence of face-likeness and region on the relationship between neural reliability and memory performance across individuals, we computed multiple linear regressions on neural reliability in the FFA and PPA in R (version 4.2.3) using the model: Pr ∼ reliability * face-likeness * ROI. Separate models were performed for each level of the localizer. As the results were comparable across localizer levels, we report only the outcome of the model on all voxels. We additionally explored the relationship between neural reliability in the searchlight cluster and memory performance using the model: Pr ∼ reliability * face-likeness.

Next, we used Bayesian linear mixed-effects models to relate trial-wise memory outcomes to neural reliability using the model: Memory ∼ reliability * face-likeness * ROI + (1 + reliability | subject) + (1 | stimulus). Memory was defined as the binary memory outcomes (hit or miss) from the old/new recognition memory task. Neural reliability was calculated within each trial as the difference in the mean similarity of a stimulus to itself across repetitions and the mean similarity of that stimulus to all other stimuli of the same face-likeness category. Separate models were computed for each level of the localizer, but we report only the outcome of the model on all voxels since all models produced comparable results. The model was replicated in the searchlight cluster without the ROI interaction.

### 2.9 Low-level feature analysis of highFL and lowFL houses

Our findings indicated differential mnemonic and representational processing of highFL and lowFL houses. Although the images were specifically selected to vary along the dimension of face-likeness, we were interested in which features of the images contributed to their face-likeness classification and whether these features alone could explain memory outcomes or neural reliability (see, e.g., Sztuka & Kühn, 2024). Here, we examined whether low-level visual features potentially contribute to differences in memory and representational similarity between highFL and lowFL houses. Visual feature decomposition was performed using a publicly-available low-level visual feature toolbox (Pohlmann et al., 2026). The following visual features were extracted from the house images: brightness, contrast, symmetry, entropy, straight edge density, non-straight edge density, and power spectrum. Visual features differing between highFL and lowFL images were determined using independent-samples *t*-tests.

As a next step, we investigated whether the features differing between highFL and lowFL houses predicted memory performance and neural reliability using Bayesian linear mixed-effects models with flat priors: Memory/reliability ∼ feature + (1 + feature | subject) + (1 | stimulus). Separate models were computed to predict reliability for the searchlight cluster as well as each level of the localizer. The localizer models predicting reliability did not converge, so reliability was averaged across participants and linear regressions were fitted: reliability ∼ feature.

## 3 Results

### 3.1 Alignment of subjective ratings of face-likeness

The current sample of participants (N = 25) rated highFL images as highly face-like on average 72.6% (SD = 19.0%) of the time. LowFL images were rated less face-like on average 80.6% (SD = 16.4%) of the time. Therefore, both the current and independent samples demonstrated strong agreement in perception of face-likeness in house images. Based on this alignment, and to ensure equal trials across conditions, we proceeded to use the independent classifications for highFL and lowFL house images.

### 3.2 Memory behavior results

We tested for differences in memory performance between highFL and lowFL houses. Dependent-samples *t*-tests revealed no differences in recognition memory performance due to face-likeness (*t*(24) = -1.13, *p* = 0.27, *d* = 0.23), but showed that lowFL houses were better-remembered than highFL houses on the forced-choice memory test (*t*(24) = -2.09, *p* = 0.047, *d* = 0.42). These findings were corroborated when houses were divided based on subjective face-likeness ratings (see Supplemental Materials). Memory performance was moderately correlated between the recognition and forced-choice tasks (*r* = 0.45, *p* = 0.025), indicating that individuals who performed better on the recognition memory task were also likely to perform well on the forced-choice memory task. As pre-registered, we used only the recognition memory results to assess subsequent brain-behavior analyses.

### 3.3 Higher representational similarity for lowFL houses in PPA

Here, we investigated whether there were regional differences in the reliability of processing highFL and lowFL houses. Analyses were conducted using ROIs of 250-voxels and 500-voxels (selected from the localizer) as well as using all voxels in the fusiform and parahippocampal gyri. Results of a 3-way ANOVA on the 250-voxel ROIs revealed an interaction between face-likeness and region (*F*(1,24) = 7.02, *p* = 0.014, *η*^2^ = 0.23), indicating that there are regional differences in the representational similarity of highFL and lowFL houses. No other main effects or interactions were significant (*p*s > 0.23). To further explore the interaction between face-likeness and region, two follow-up ANOVAs were conducted separately for the FFA and PPA. The ANOVA in the PPA revealed a main effect of face-likeness (*F*(1,24) = 4.45, *p* = 0.045, *η*^2^ = 0.16), indicating that activation patterns of lowFL houses (*M* = 0.043, *SD* = 0.038) were more similar in the PPA than activation patterns of highFL houses (*M* = 0.035, *SD* = 0.034). No other main effects or interactions were significant, either in the PPA or FFA (*p*s > 0.45).

Results of the ANOVAs on the 500-voxel ROIs qualitatively concurred with the results in the 250-voxel ROIs, except for an additional main effect of ROI in the 3-way ANOVA (*F*(1,24) = 9.44, *p* = 0.005). For the full statistical report, refer to the supplements.

Results of the 3-way ANOVA using all voxels showed a marginal 3-way interaction (*F*(1,24) = 3.94, *p* = 0.058, *η*^2^ = 0.14) as well as a marginal interaction between similarity and region (*F*(1,24) = 3.88, *p* = 0.060, *η*^2^ = 0.14; see Figure 4). A follow-up ANOVA in the FFA revealed a main effect of face-likeness (*F*(1,24) = 6.33, *p* = 0.019, *η*^2^ = 0.21), in which lowFL houses (*M* = 0.15, *SD* = 0.076) were represented more similarly in the FFA than highFL houses (*M* = 0.14, *SD* = 0.066). The follow-up ANOVA in the PPA demonstrated a main effect of similarity (*F*(1,24) = 6.45, *p* = 0.018, *η*^2^ = 0.21) as well as a main effect of face-likeness (*F*(1,24) = 7.54, *p* = 0.011, *η*^2^ = 0.24). Between-stimulus similarity (*M* = 0.027, *SD* = 0.021) was greater than within-stimulus similarity (*M* = 0.021, *SD* = 0.028). LowFL houses (*M* = 0.029, *SD* = 0.027) were represented more similarly than highFL houses (*M* = 0.020, *SD* = 0.022). Furthermore, there was a marginal interaction between reliability and face-likeness (*F*(1,24) = 4.12, *p* = 0.054, *η*^2^ = 0.15), in which reliability for lowFL houses was zero (*M* = 0.00, *SD* = 0.018), but reliability for highFL houses was negative (*M* = -0.012, *SD* = 0.021). Therefore, these findings suggest that the PPA did not demonstrate representational reliability as per the definition of having higher within-stimulus similarity than between-stimulus similarity.

**Figure 4.**
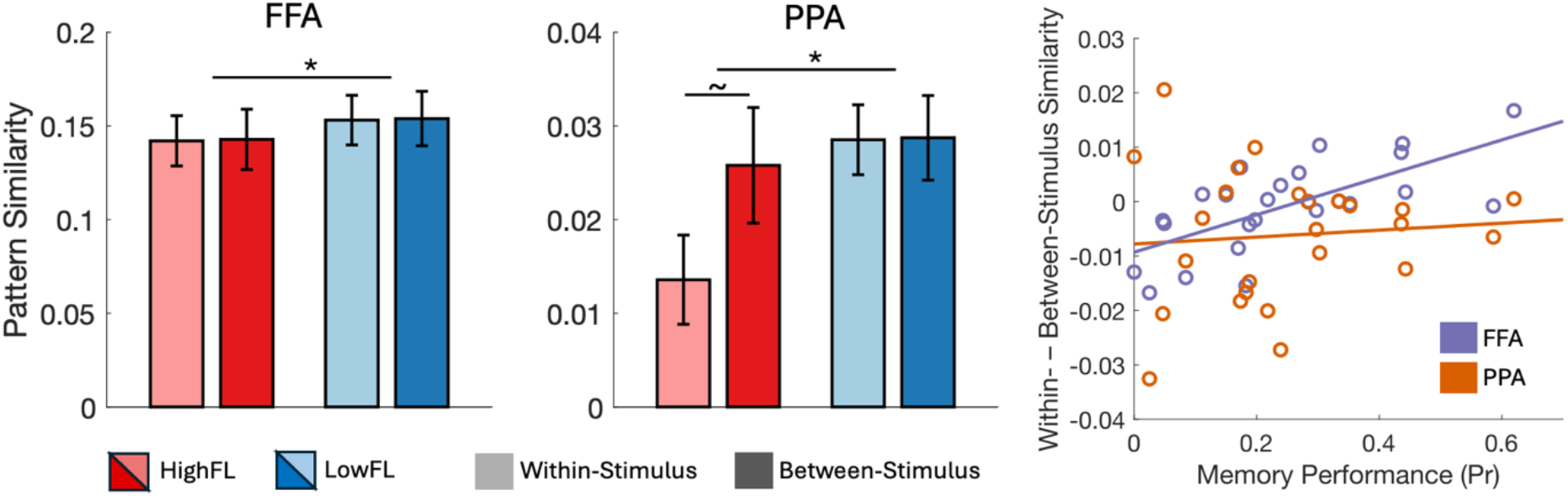
Pattern similarity in the FFA (left) and PPA (middle) across all voxels in the anatomical regions (note that the y-axis differs between the ROIs). LowFL houses (blues) were represented more similarly than highFL houses (reds) in both ROIs. Between-stimulus similarity (darker colors) tended to be greater than within-stimulus similarity (lighter colors) in the PPA, particularly for highFL houses. Asterisks indicate significant effects (*p* < 0.05) and the tilde represents a marginally significant effect. The relationship between neural reliability (the difference in within- and between-stimulus similarity) and memory performance (right). Higher neural reliability was associated with better memory performance across individuals, particularly in the FFA (purple) relative to the PPA (orange).

Together, our findings consistently demonstrated that lowFL houses were represented more similarly than highFL houses in the PPA. However, we did not observe that face-likeness modulated representations in the FFA as we expected. This finding was validated by several exploratory assessments (see Supplemental Materials), all of which failed to produce a significant effect of face-likeness in the FFA. Furthermore, we did not find that within-stimulus similarity was greater than between-stimulus similarity, which would be indicative of neural reliability. Therefore, we expanded our search for neural reliability by running a whole-brain searchlight similarity analysis.

### 3.4 Higher neural specificity for lowFL houses in early visual cortices

We were particularly interested in how face-likeness modulates representational reliability in the FFA. However, in the previous analysis, we did not find evidence of representational reliability in either the FFA or PPA. Here, we searched the whole brain for differences in neural reliability between highFL and lowFL houses. Our searchlight resulted in one cluster in which lowFL houses were represented with greater reliability than highFL houses (see Figure 5). This cluster spanned parts of the bilateral calcarine cortex, lingual gyrus, superior occipital gyrus, and cuneus (*k* = 1833 voxels).

**Figure 5.**
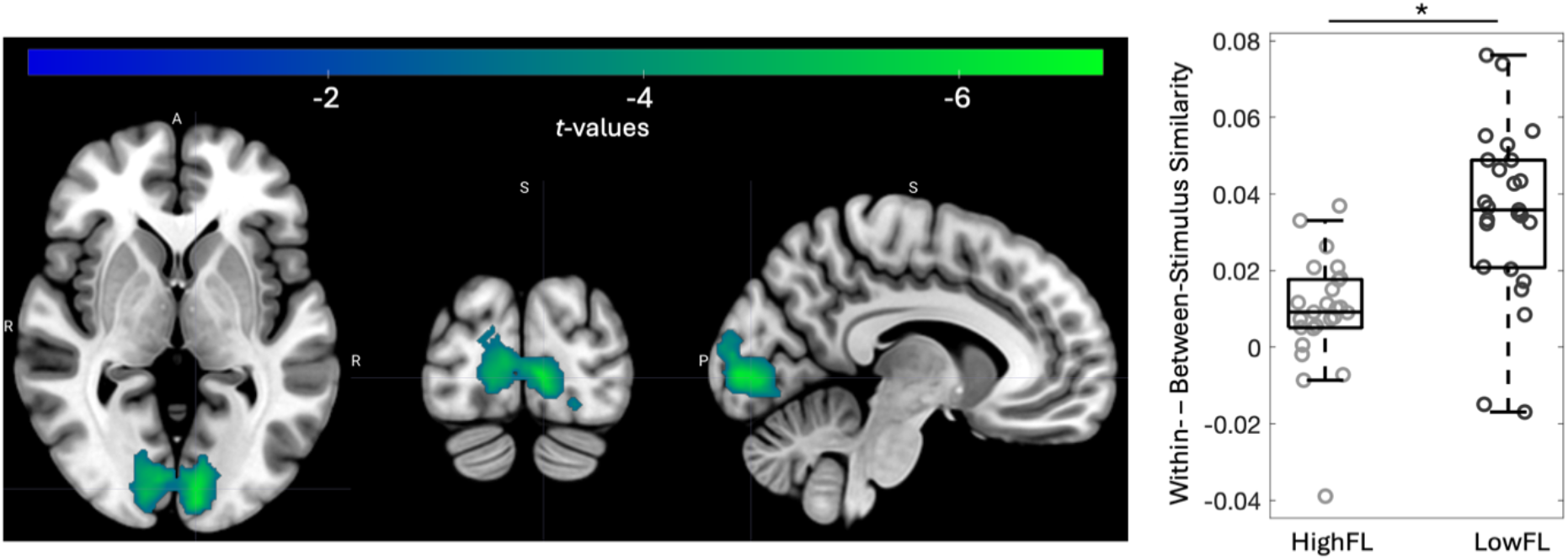
Searchlight cluster (left) revealing regions in which lowFL houses are represented more reliably than highFL houses. Box plots (right) show the distribution of neural reliability for highFL (light gray) and lowFL (dark gray) houses in the searchlight cluster.

Follow-up analyses on this cluster indicated that neural reliability for both lowFL and highFL houses was significantly greater than zero (lowFL: *t*(24) = 7.77, *p* < 0.001; highFL: *t*(24) = 3.13, *p* = 0.005).

### 3.5 Neural reliability predicts interindividual differences in memory performance

Results of a multiple linear regression to predict memory performance revealed a marginally significant model (*F*(7,92) = 1.97, *p* = 0.068, *R*^*2*^ = 0.13). Individual predictors demonstrated a main effect of neural reliability (*t* = 2.81, *p* = 0.006), indicating that individuals with higher neural reliability tend to have better overall memory performance. Furthermore, there was an interaction between neural reliability and ROI (*t* = -2.39, *p* = 0.019), suggesting that the strength of the relationship between reliability and memory is attenuated in the PPA compared with the FFA. No other main effects or interactions reached significance (*p*s > 0.30). The multiple linear regression to predict memory performance from reliability in the searchlight cluster did not reveal a significant model (*F*(3,46) = 1.84, *p* = 0.15, *R*^*2*^ = 0.11).

Trial-wise analyses relating neural reliability to memory outcomes did not demonstrate any main effects or interactions in the ventral visual regions nor in the searchlight cluster (all 90% and 95% CIs included 0).

### 3.6 Visual feature variability between highFL and lowFL houses

Here, we explored whether inherent differences in low-level visual features between highFL and lowFL houses could explain the observed mnemonic and representational differences between the stimulus categories. Results of *t*-tests demonstrated differences in symmetry (*t*(38) = 5.00, *p* < 0.001) between highFL and lowFL houses, revealing that symmetry was greater in highFL houses (*M* = 0.58, *SD* = 0.06) than lowFL houses (*M* = 0.48, *SD* = 0.06). Brightness (*t*(38) = -1.06, *p* = 0.30), contrast (*t*(38) = 0.17, *p* = 0.87), entropy (*t*(38) = -1.30, *p* = 0.20), straight edge density (*t*(38) = 1.36, *p* = 0.18), non-straight edge density (*t*(38) = -1.58, *p* = 0.12), and power spectrum (*t*(38) = -1.17, *p* = 0.25) did not differ between highFL and lowFL houses.

Linear mixed effects modeling demonstrated that memory was inversely related to symmetry (est. = -0.68, 95% CI: [-1.16, -0.20]), indicating that less-symmetrical houses tended to be better remembered than highly symmetrical houses. Furthermore, the interaction between symmetry and face-likeness was significant (est. = 0.85, 95% CI: [0.21, 1.50]), suggesting that less-symmetrical lowFL houses are better-remembered than more-symmetrical lowFL houses. However, only face-likeness, not symmetry or the interaction, significantly predicted representational reliability in early visual cortices (est. = -0.04, 95% CI: [-0.07, -0.01]). Since the linear mixed effects models predicting representational reliability in the localizer ROIs did not converge, we used linear regression to relate symmetry to reliability in these regions. However, no main effects reached significance (*p*s > 0.16). 19

## 4 Discussion

The main objective of this study was to evaluate the relationship between representational reliability and memory performance in ventral visual cortices which preferentially process face and house stimulus categories (i.e., FFA and PPA, respectively). Crucially, we were interested in controlling inherent stimulus differences between face and house stimuli to disentangle potential regional differences in the mnemonic function of the corresponding neural representations. We achieved this manipulation by assessing a specific set of house stimuli in which half had face-like characteristics and the other half did not. However, contrary to our expectations, we did not find evidence for face-specific processing in the FFA in response to the face-like houses. Nevertheless, we uncovered several interesting findings warranting discussion. First, we found that less face-like houses tended to be more memorable than face-like houses. Furthermore, less face-like houses elicited more consistent neural representations in early visual cortices, but not in ventral visual cortices, across repeated presentations compared with face-like houses. Finally, we discuss the potential role of symmetry in explaining mnemonic differences between face-like and non-face-like houses.

Prior research has indicated that memory performance scales with face-likeness, with faces or non-face stimuli with face-like characteristics being more memorable than buildings or stimuli with few face-like qualities (Brady et al., 2019; Kapsetaki & Zeki, 2022). This selective memorability for faces has been proposed to reflect familiarity (i.e., more expertise or more self-similarity), preference, or inherent stimulus properties specific to faces (e.g., symmetry; Kapsetaki & Zeki, 2022). Consequently, we would have expected more face-like houses to be better-remembered than less face-like houses. However, we did not find evidence confirming that face-like houses were more memorable than less face-like houses, and, contrary to the existing evidence, we found a tendency that less face-like houses were more memorable (see also Supplementary Materials). This contradictory finding was further supported by evidence suggesting that less-symmetrical houses were more memorable than more-symmetrical houses. One possibility is that symmetry supports memory for faces due to the fact that symmetry is a prototypical trait of faces. In contrast, symmetry may maladaptively influence memory for houses because symmetry constrains the variety of features that may stand out in one’s memory since symmetrical images tend to contain less information (due to mirroring) than nonsymmetrical images. Our finding that reduced symmetry was beneficial for memory of lowFL houses supports this interpretation. One recent study found that symmetry enhanced performance on a working memory task (Sztuka & Kühn, 2025). Together, these findings may suggest that symmetry supports mnemonic processing across short time spans, while impairing long-term memory outcomes. Future studies might examine how stimulus features and their prototypicality influence memorability of different stimulus categories.

Our findings point to stimulus-specific reliability of neural representations in early visual cortices, but not in ventral visual cortices. Visual representations are assumed to be hierarchically organized with early visual regions representing individual stimulus properties (Kamitani & Tong, 2005) and ventral visual regions representing entire stimuli and stimulus categories (Collins & Olson, 2014; Cowell et al., 2010; Epstein & Kanwisher, 1998; Kanwisher et al., 1997). Cognitive processing may influence the stability of neural representations in higher-order regions, whereas low-level visual cortices may remain relatively unaffected by cognition and thus more reliably represent stimulus features (Collins & Olson, 2014). Therefore, the finding of reliability in the calcarine cortex and surrounding areas, but not in ventral visual regions, may not be so surprising. Interestingly, stimulus-specific reliability in early visual cortices was modulated by face-likeness, in which lowFL houses were represented more reliably than highFL houses. This effect may reflect low-level feature differences between lowFL and highFL houses, with some features characteristic to lowFL houses being more stably represented than others. However, we only found one feature (symmetry) to significantly differ between highFL and lowFL houses, and symmetry could not explain reliability in the early visual cortical cluster. Our finding aligns with prior evidence showing that neural activity patterns in V1 and V2 do not differ between symmetrical and asymmetrical dot patterns (Van Meel et al., 2019). Together, low-level visual features likely do not explain why lowFL houses are represented more stably in early visual cortices than highFL houses. However, to our knowledge, this is the first investigation into whether stimulus properties influence reliability of neural representations, so future studies are needed to corroborate these findings.

The premise of this study was to evaluate whether neural reliability relates to memory performance differentially in FFA and PPA. The role of stable neural representations in supporting mnemonic processing has been previously demonstrated as a mechanism both to explain interindividual variability in memory performance (Koen et al., 2019; Pauley et al., 2023) as well as to understand why some events are remembered and others forgotten (Hasinski & Sederberg, 2016; Xue et al., 2010). Although we do not provide strong corroborating evidence, we attribute this to the relatively low sample size and low number of trials in the current study rather than questioning the validity of previous findings. Furthermore, we contribute a marginal effect showing that individuals with more stable neural representations in ventral visual cortices also tend to have overall better mnemonic abilities. While evidence converges implicating PPA neural representations in memory outcomes, the mnemonic function of neural representations in the FFA is more contentious. We predicted that inherent stimulus differences between faces and houses may explain these apparent regional differences in memory function. Although our manipulation was not effective in producing face-like responses in the FFA for face-like houses, our results nevertheless lent support for a stronger link between memory and representational reliability in the FFA than the PPA. Therefore, stimulus properties differing between faces and houses may indeed influence memory-related representations in ventral visual cortices, though more research is needed to validate our findings.

The findings of the present study have further implications for research on the neural processing of built environments and related fields such as neuroarchitecture. Prior fMRI research on architecture has largely focused on aesthetic and subjective responses (Vartanian et al., 2013, 2015, 2024), as well as neural correlates of perceived qualities like fascination and hominess (Coburn et al., 2020). While there is some evidence that higher-level architectural features may be represented in regions like the FFA (Choo et al., 2017), these studies provide limited insight into how specific perceptual features of buildings relate to memory and neural representations. Recent evidence implicates engagement of the hippocampus and PPA in memorable architectural experiences (Gregorians et al., 2026). Our findings that less symmetrical house facades are more memorable and have more reliable neural representations than symmetrical facades could inform future experimental or applied work in urban and architectural contexts.

We would like to point out two limitations of the present study. First, since faces were not presented in the main fMRI task, we cannot directly compare the reliability of face representations in the FFA to the house representations. A direct comparison in which the across-voxel patterns of face stimuli are related to the highFL and lowFL conditions could provide a further index to see if the neural activity patterns are face-like. In lieu of a direct comparison, we assume that voxels preferentially activated in response to face stimuli would also most reliably represent face stimuli across repetitions (as shown in previous studies, e.g., Hasinski & Sederberg, 2016). Face stimuli would additionally lend support for our memory analyses to understand if memory performance differed between faces and face-like houses and the role of symmetry in memory for faces. A second limitation is that the face-like houses were evidently not sufficiently “face-like” to elicit face-specific activity patterns in the FFA. The distribution of subjective ratings of face-likeness (see Supplemental Materials) further suggest that participants did not perceive clear face-like qualities in the houses. One possibility is that the DalHouses stimulus set, acquired in Canada, were relatively unfamiliar to German participants and did not comparably engage perception of face-likeness. However, a recent intercultural investigation of the DalHouses indicates minimal cultural differences in anthropomorphism of these exact stimuli (Weber et al., 2024), therefore we believe this to be an unlikely explanation. Rather, the subjective ratings indicate that face-likeness is predominant in just a few houses, not across all stimuli in the highFL condition. Further, participants were not cued to attend face-likeness before the scanning session. Thus, they may not have noticed the implicit modulation of face-likeness in the houses (but, see Kühn et al., 2014, who showed FFA activation to highly face-like cars without explicit cueing). Future studies may consider assembling a set of face-like houses with more consistent face-like characteristics or consider priming participants to attend face-like characteristics in non-face stimuli.

Our results indicate that face-likeness of house stimuli modulates mnemonic outcomes, representational reliability, as well as their interrelatedness. In particular, we provide preliminary evidence that symmetry may be a key perceptual factor underlying these effects, potentially shaping mnemonic outcomes. Together, we contribute a novel analysis aimed at understanding the mnemonic impact of reliable neural representations.

## Data availability

The behavioral data and code will be made available on OSF at the following link: https://osf.io/23tvb/overview. The brain data is available upon request to the corresponding author.

## Author contribution

**Claire Pauley**: Conceptualization, Data curation, Formal analysis, Investigation, Software, Visualization, Writing - Original Draft. **Izabela Maria Sztuka**: Conceptualization, Data curation, Investigation, Methodology, Project administration, Software, Writing – Review & Editing. **Nour Tawil**: Conceptualization, Investigation, Resources, Writing – Review & Editing. **Simone Kühn**: Conceptualization, Supervision, Writing – Review & Editing. 3

## Acknowledgments

This work was conducted at the Max Planck Dahlem Campus of Cognition (MPDCC) of the Max Planck Institute for Human Development, Berlin, Germany. We would like to especially thank Sonali Beckmann and Nadine Taube for their assistance in MR data acquisition as well as to Jiaona Hu for support in data collection.

