## Supplements for "Face in the facade: How face-likeness modulates memory and neural representations"

*Memory performance between subjectively-rated highFL and lowFL houses*

We re-tested for differences in memory performance between highFL and lowFL houses as determined by individuals' subjective ratings of face-likeness. Dependent-samples *t*-tests revealed marginal differences in recognition memory performance due to face-likeness ($t(24) = -2.03$, $p = 0.054$), as well as significant differences in forced-choice memory performance ($t(24) = -2.73$, $p = 0.012$). In both cases, lowFL houses were more likely to be remembered than highFL houses.

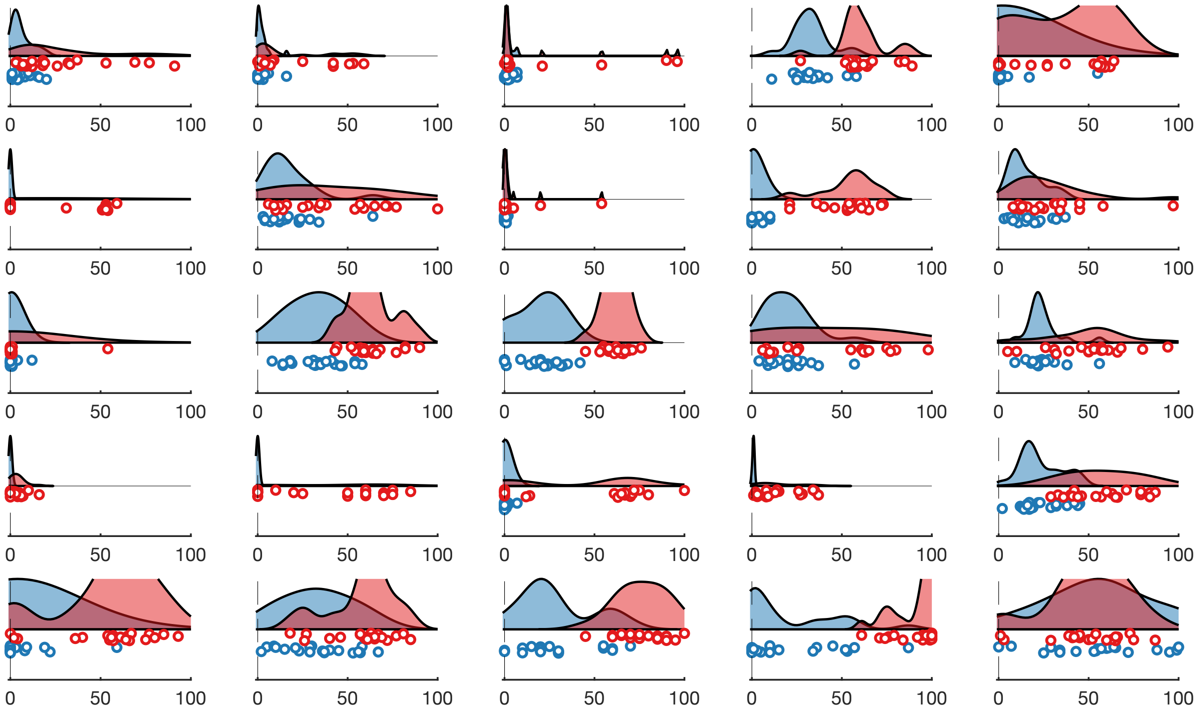


*Figure S1.* Subjective ratings of face-likeness for each participant (25 participants in total). Participants rated their subjective face-likeness to each individual stimulus on a scale of 0 (not at all face-like) to 100 (highly face-like). Blue indicates houses independently rated as less face-like, while red indicates houses independently rated as highly face-like. Responses to individual stimuli are plotted below violin plots.

*Complete report of ANOVAs in 500-voxel ROIs*

Results of the 3-way ANOVA on the 500-voxel ROIs showed a main effect of ROI ($F(1,24) = 9.44$, $p = 0.005$) as well as a marginally significant interaction between face-likeness and region ($F(1,24) = 4.15$, $p = 0.053$). The 2-way ANOVA in the PPA demonstrated a main effect of face-likeness ($F(1,24) = 5.77$, $p = 0.024$), indicating that activation patterns of lowFL houses ($M$ = 0.038, $SD$ = 0.029) were more similar in the PPA than activation patterns of highFL houses ($M$ = 0.029, $SD$ = 0.026). No other main effects or interactions reached significance in any of the models ($p$s > 0.21).

*Mean activation is higher for lowFL houses than highFL houses in PPA*

We investigated whether mean activation differed between highFL and lowFL houses in the FFA and PPA. The activation maps were generated using an event-related GLM in which regressors modeled the highFL houses, lowFL houses, fixation crosses, as well as six motion parameters. High- and lowFL conditions were contrasted to the fixation crosses, resulting in whole-brain $t$ maps of highFL and lowFL activation. The mean activation across voxels in each of the participant-specific masks as well as the whole anatomical ROIs were evaluated separately in 2 (highFL/lowFL) x 2 (FFA/PPA) ANOVAs and follow-up dependent-samples $t$-tests.

For all masking levels, we found an interaction between face-likeness and region ($p$s < 0.026, indicating that face-likeness differentially modulated activation in the FFA and PPA. For the 250- and 500-voxel levels, there was higher activation in response to lowFL houses than highFL houses in the PPA ($p$s < 0.004), but no differences in activation in the FFA ($p$s > 0.37). In contrast, when assessing the entire anatomical regions, the FFA showed higher activation in response to the lowFL houses than highFL houses ($t(24) = -3.34$, $p = 0.003$), while the FFA showed no differences in activation ($t(24) = -1.63$, $p = 0.12$).

*Hemisphere differences between highFL and lowFL houses in FFA and PPA*

Here, we investigated whether the left and right hemispheres differentially processed face-likeness potentially masking any effects of neural specificity in the bilateral FFA and PPA. As such, we performed 2 3-way ANOVAs (within-stimulus/between-stimulus x highFL/lowFL x left/right hemisphere): one in the FFA and one in the PPA. The participant-specific top 250 and 500 voxels were divided based on the hemisphere. Participants with fewer than 10 voxels in one hemisphere were excluded from the analyses, resulting in one exclusion in the 250-voxel FFA ANOVA.

In the 250-voxel FFA ANOVA, there was a 3-way interaction between specificity, face-likeness, and hemisphere ($F(1,23) = 6.31$, $p = 0.020$). No other interactions or main effects reached significance ($p$s > 0.12). Follow-up ANOVAs in each hemisphere indicated no significant main effects or interactions for specificity or face-likeness ($p$s > 0.081).

No main effect of hemisphere or interactions with hemisphere reached significance in any other model ($p$s > 0.13). We therefore concluded that neural specificity was not found in individual hemispheres.

*Evaluating effects of face-likeness in clusters of adjacent voxels in the FFA*

In this analysis, we were interested if neural specificity would be detected in the FFA when clusters of adjacent voxels were considered (rather than the most-selective 250 or 500 voxels regardless of spatial proximity). To this end, we implemented a cluster-based approach in which adjacent voxels exceeding an uncorrected threshold of $p < 0.005$ were defined as a cluster. For each participant, the cluster with the highest average t-value in the faces > houses localizer contrast was considered within the bilateral fusiform gyrus mask. Eight participants did not have adjacent voxels reaching this threshold and were excluded from the present analysis. The same pattern similarity analysis and ANOVA were performed using these cluster-based voxels. A 2 (within-stimulus/between-stimulus) x 2 (highFL/lowFL) ANOVA revealed no main effects or interaction ($p$s > 0.48).

*Considering interindividual differences in anthropomorphism*

Some individuals have a stronger tendency to anthropomorphize, which has been associated with greater activation in the FFA (Kühn et al., 2014). Therefore, we additionally explored whether isolating individuals with higher anthropomorphic tendencies would uncover differences in neural specificity between high- and lowFL houses. During the post-scanner behavioral session, participants completed the Individual Differences in Anthropomorphism Questionnaire (IDAQ; Waytz et al., 2010). Anthropomorphism was scored as the sum of responses in the IDAQ, with higher scores reflecting greater anthropomorphic tendencies ($M$ = 34.48, $SD$ = 12.15). We repeated the representational similarity analyses (methods described in Section 2.6) including only individuals who scored higher than the mean ($N$ = 14). The results agreed qualitatively with the findings reported in Section 3.3, indicating no significant differences in neural specificity between high- and lowFL houses, even in individuals with anthropomorphic tendencies.

*Relating representational similarity to interindividual differences in memory performance*

Our results in Section 3.3 consistently indicated that lowFL houses were represented more similarly than highFL houses. Here, we explored whether representational similarity predicted memory performance and whether this was modulated by face-likeness or region. Linear mixed effects models were used to predict memory performance from neural specificity (i.e., Pr ~ similarity * face-likeness * ROI) for each level of the localizer. However, no models reached significance ($p$s > 0.24), indicating that overall representational similarity in the FFA and PPA was not related to memory ability in our sample.
